# White and Clearing: New Optical Tissue Clearing Method

**DOI:** 10.1101/2025.01.30.635653

**Authors:** Vicente Llorente, Elena Fernández-Cortés, Daniel Sanderson, Julia Salazar, Jorge Ripoll, Manuel Desco, María Victoria Gómez-Gaviro

## Abstract

Optical tissue clearing techniques can render large-scale tissue samples and whole organs transparent to apply cell-resolution 3D imaging techniques, revolutionizing the fields of biological research and pathological diagnosis, and providing never-before seen structural information. There are several widely used protocols, but they have shortcomings, both in complexity of use and in suboptimal results in certain tissues and organs.

Here, we describe a new method dubbed “White and Clear” (WaC) using 2,2’-thiodiethanol (TDE) as a dynamic Refractive Index (RI)-matching reagent alongside a lipid-extraction and a pigment-bleaching step using dilute hydrogen peroxide (H2O2), to provide ease of use and enhanced clearing effect on all kinds of samples and allow for the clearing parameters to be finely adjusted in order to obtain the best results in each situation.

## 1. Introduction

The dynamics of biological tissues and organs are heavily affected by their structure, as they exist, develop, and interact with each other in a three-dimensional space. Any information concerning this structural disposition could be key in deciphering the physiology of any biological process, be they natural or the result of some pathology or condition.

The microscopic study of biological samples has been one of the main fields of biology and medicine for a long time, but the methods it employs usually rely on the sectioning of samples into slices, thin enough to allow penetration of light without being absorbed or scattered, thus enabling the use of microscopy techniques. This, however, results in the loss of most three-dimensional information, as samples are physically cut and artifacts are introduced, greatly hindering the recovery of such an important source of information.

To solve this, optical tissue clearing techniques were introduced to render large-scale biological samples transparent and increase light penetration, thus enabling the use of microscopy techniques to acquire whole samples while keeping their structural information intact and ripe for analysis. These methods were described more than a century ago (Spalteholz 1914) but have become truly popular after the development of high-resolution three-dimensional microscopy techniques actually capable of taking advantage of these effects, such as two-photon, confocal or selective plane illumination microscopy (SPIM) (Gómez-Gaviro et al. 2020).

While the original approaches were based on tissue dehydration and solvent-based clearing, more recent ones explored the field of aqueous-based reagents, which offer advantages over the former; most notably, the relatively low toxicity of its components compared to solvent-based methods, as well as the reduction in damage to samples and fluorescence quenching (Richardson and Lichtman 2015). The core principles of their action are the same as the older methods: removal of tints and pigments, extraction of lipids, and homogenization of Refractive Index (RI) values through the sample, to prevent light from being absorbed or scattered haphazardly (Gómez-Gaviro et al. 2020; Tainaka et al. 2018). While originally developed for specific tissues and organs in particular, these have expanded to cover a wider variety through trial and error as well as exhaustive optimizations. Some examples of this kind of use in tissues other than the original target ones (in this case, murine brain) include murine colon (Zufiria et al. 2016), chicken embryo (Bocancea et al., n.d.; Goḿez-Gaviro et al. 2017), murine heart (Nehrhoff et al. 2016; 2017), whole mice and rat (using perfusion-based methods for clearing reagent application) (Tainaka et al. 2014; Matryba et al. 2018), and, more recently, human melanoma and melanoma xenograft model (Llorente et al. 2022).

A relatively recent arrival to the world of aqueous-based tissue clearing, 2,2’-thiodiethanol (TDE), is an organosulfur compound with both polar and nonpolar properties, used as a solvent and as a chemical building block in industrial applications. TDE was described as a perfect candidate for use as mounting media in microscopy, as it presents a very high RI value (∼1.515) which can be easily brought down to any value up to 1.31 by simply diluting it in water or any other similar liquid (Staudt et al. 2007).

This property also shows promise for the purposes of tissue clearing, as the components of most tissues display high RI values (1.41-1.44), and the ability to easily make fine adjustments to RI would be helpful for RI-homogenization steps. And yet, all attempts to use TDE for tissue clearing have provided lukewarm results so far, lacking suitability for use in full-scale organs and large-scale samples (even if it was reasonably efficient on smaller samples) (Aoyagi et al. 2015), or being applied in combination with existing methods, with the ensuing complexity (Costantini et al. 2015). There has been certain success in larger samples belonging to some specific tissues (Sun et al. 2022), although this is mostly achieved on a case-by-case basis.

The presence of pigments in tissues and organs is an important source of opacity, as these substances absorb light on a wide range of wavelengths and thus prevent its further penetration into the sample even if other conditions would allow for it. Pigments are also a problem in conventional 2D-imaging techniques, as for example they can interfere with the dyes used in chromogenic immunohistochemistry to label target proteins and thus prevent proper analysis of histopathology samples. This issue is particularly egregious in melanoma samples, as they present overwhelmingly high concentrations of melanin. However, there are other cumbersome pigments found in healthy internal organs.

Most tissue opacity, along with color, is removed upon the standard delipidation step customarily included in solvent- and detergent- based tissue clearing protocols. While many samples respond well to lipid removal-induced refraction index matching and become translucent, some organs - such as the liver or spleen - may prove more problematic and retain stubborn layers of opaque pigment. Therefore, an in-depth understanding of the biological pigments dyeing internal structures is of utmost importance for the appropriate optimization of the currently available clearing methods.

Observable color arises, mainly, from two different optical phenomena: interference and scattered light. The former is the very basis of “true” pigments which, unlike scattered colors, remain unchanged by crushing or otherwise mechanical disruption. Although both may act in tandem to produce a wide array of hues, interference-borne pigmentation is the more relevant source of color and the larger hurdle to surpass as far as organ clearing is concerned.

Opacity, rather than color itself, is another issue entirely, as light scattering happens to be the principal culprit. Tissues are universally heterogeneous structures, and this inhomogeneity is caused by diverse inter- and intra-cellular components, wherein membrane lipids are among the most disruptive (Tuchin 2016) and, historically, a target for removal in organ clearing (Richardson and Lichtman 2015). The main goal of current clearing methods, rather than complete lipid removal, is the matching of refraction index across as many histological structures as possible, for which delipidation still proves effective in solvent-, aqueous- and detergent-based protocols. However, non-lipophilic pigments may remain unchanged and unaccounted for.

In literature vertebrate biological pigments are largely grouped into carotenoids (subdivided into xanthophylls and carotenes, the latter of which are more abundant and relevant to our discussion), hemoproteins, lipochromes, melanin, and others (Piña-Oviedo, Ortiz-Hidalgo, and Ayala 2017). As lipophilic structures, carotenes are associated with lipid rich tissues and are understood to bind to intercellular lipids (Choe et al. 2019). Interestingly, they are also soluble in several fat solvents (Su, Rowley, and Balazs 2002), making it so that the previously mentioned delipidation step rids organs not only of the opacity conferred by light-blocking fat, but their orange/yellow pigmentation as well. Cells themselves remain intact as they do not absorb any wavelength of visible light (Inyushin et al. 2019). Thus, organ clearing protocols that rid tissues of their inherent pigments render structures translucent rather than white.

Matters get trickier regarding the color red. In the same manner that plants are ubiquitously green due to the essential role of chlorophyll in electron transfer via photosynthesis, an analogous process involving a biological pigment also occurs in vertebrates: cellular respiration through oxygen transport and storage via hemoproteins, which dye innards red. Stating that heme groups are the most versatile protein cofactors is far from hyperbolic; within the realm of the mammalian body, hemoprotein functions span well beyond the scope of gaseous molecule delivery and are an essential component in redox sensing (Gilles-Gonzalez and Gonzalez 2005), catalysis (Munro, Girvan, and McLean 2007), gene regulation (Zhang and Guarente 1995; Faller et al. 2007) and — simultaneously bestowing a deep shade of red to mitochondria-rich organs — electron transfer reactions (Nicholls 1996) facilitated by cell membrane associated cytochromes.

One of CUBIC’s standard components, N,N,N,N-tetrakis(2- hydroxypropyl)ethylenediamine, an aminoalcohol, is theorized to aid in hemoprotein removal through heme dissociation, although this effect was not intended during the development of the method and is largely incidental (Susaki et al. 2015). Its depigmentation prowess, however, is low (Ren et al. 2021) and calls for the introduction of a decolorizing agent able to remove the hemoproteins responsible for the light-absorbing red that hinders tissue clearing.

To sidestep this issue, bleaching techniques have been developed to remove pigments from samples while avoiding damage to samples and target proteins (or at least reducing it to an acceptable minimum). A combination of potassium permanganate and oxalic acid is one of the most common methods, and, while effective enough, it is known to cause damage to certain antigens, thus preventing them from being properly labeled and studied (Foss et al. 1995; Guy Edward Orchard and Clone 1998; G. E. Orchard 1999). On the other hand, the use of dilute hydrogen peroxide (H2O2), applied at high temperatures to boost its bleaching action, offers a simpler and faster alternative that also better preserves tissue structure and immunogenicity (Li et al. 1999; Momose, Ota, and Hayama 2011; Liu et al. 2013; Chung et al. 2016).

As bleaching with dilute H2O2 has been stated as a quick, easy, and effective method to remove pigments from small-size tissues and biopsy samples, we wanted to test whether this protocol could be applied to whole organs and thick tissue samples to improve the pigment bleaching effect of the clearing protocol and reduce light absorption as a result. Likewise, the potential shown by TDE for its inclusion in optical tissue clearing protocol moved us to study it as a potential replacement for RI-homogenizing reagents or steps in standard aqueous-based tissue clearing methods, in order to sidestep the downsides or shortcomings of existing techniques (such as difficulty of manipulation and preparation) and streamline the process of successfully clearing whole organs and large-scale samples for biological study.

## 2. Materials and Methods

### 2.1 Mice

All the experimental procedures were approved by the hospital’s Ethics Committee for Animal Research (*Comisión de Ética de Experimentación Animal, CEEA*) and by the Comunidad Áutonoma de Madrid (CAM) with PROEX 269.1/22 and were carried out in full compliance with ARRIVE guidelines.

C57BL/6 mice, 8–10-weeks-old, weighing 25–30 g, housed in an air-conditioned room with a 12 h light/dark cycle and free access to water and chow were used in this study.

Adult male LysMcre^+/−^, mT/mG and AlbCre+/−, mT/mG mice were also used in this study and maintained in similar conditions. LysMcre+/−, mT/mG mice express membrane-targeted tandem dimer (Td) Tomato (a red fluorescent protein) in all cells except for those with a myeloid cell lineage, which instead express membrane-targeted enhanced green fluorescent protein (EGFP). AlbCre+/−, mT/mG mice express membrane-targeted tandem dimer (Td) Tomato (a red fluorescent protein) in all cells except for those with a hepatic cell lineage, which instead express membrane-targeted enhanced green fluorescent protein (EGFP). Further information can be found in (Nehrhoff et al. 2016; Muzumdar et al. 2007).

### 2.2 Sample processing

Animals were sacrificed by transcardial perfusion. Prior to perfusion, a ketamine/xylazine mixture was administered via intraperitoneal injection to deeply anesthetize the animals. The heart was perfused with 15 ml of PBS 1× to flush blood out of the animal’s vessels, followed by 40 ml of 4% PFA in PBS 1× for fixation.

The organs were excised and incubated in PFA 4% for 24h at 4°C. Stomach and intestines were manually cleared of partially digested food whenever possible to avoid their interference in the clearing process. Samples were then rinsed with PBS and stored in Sodium Azide 0.02% in PBS 1× at 4°C for preservation until their further analysis.

For the modified perfusion-based clearing protocol, the standard perfusion protocol was applied, followed by perfusion with 40 mL of Reagent 1 (R1) and 10 mL of 10% w/v hydrogen peroxide (H2O2) solution (where applicable). Samples were then immersed in R1.

### 2.3 Tissue clearing and immunohistochemistry

#### 2.3.1 Standard tissue clearing reagents

The standard tissue clearing protocol is composed of two separate steps with their corresponding reagents: the first reagent (known as R1, or Reagent 1) removes lipids and pigments and hyper-hydrates the samples, and the second (known as R2 or Reagent 2), homogenizes the RI values throughout the sample. RI values only remain homogeneous for as long as the sample is kept in R2, and immersion in any other reagents reverts transparency, requiring the bleaching and immunohistochemistry-immunofluorescence (IHC-IF) procedures to be carried out between R1 and R2. R1 and R2 were prepared according to the original protocol; a summary of their compositions can be found in **Table 1**.

**Table 1:** Composition of Reagent 1 (R1) and Reagent 2 (R2) from the standard optical tissue clearing protocol.

| COMPONENT | Reagent 1 (R1) | Reagent 2 (R2) |
| --- | --- | --- |
| Distilled water | 40% w/v | 15% w/v |
| N,N,N,N-tetrakis(2-hydroxypropyl)ethylenediamine | 20% w/v | N/A |
| Urea | 25% w/v | 25% w/v |
| Triton X-100 | 15% w/v | N/A |
| Sucrose | N/A | 50% w/v |
| Triethanolamine | N/A | 10% w/v |

#### 2.3.2 Bleaching solution

The hydrogen peroxide (H2O2)-based bleaching solution was prepared as reported to enable high bleaching efficiency and minimize damage to tissue morphology: 10% w/v concentration in PBS 1× and incubation at 65°C in glass containers (Liu et al. 2013). H2O2 30% w/v (Foret, Peroxfarma, Barcelona, Spain) was diluted with PBS to create a 10% w/v solution.

#### 2.3.3 2,2’-thiodiethanol (TDE)

The 2,2’-thiodiethanol (TDE) clearing solution is a RIR, and as such, acts by homogenizing the RI values of the sample, and its effects are reverted once the sample is transferred to any other solution; thus, bleaching and IHC-IF procedures for SPIM 3D imaging were also carried out before incubation in TDE.

≥99% TDE (Sigma Aldrich, Merck KGaA, Darmstadt, Germany) was diluted with PBS 1× to create solutions of different concentrations: 70% v/w, 80% v/w, 90% v/w and 97% v/w.

#### 2.3.4 Immunohistochemistry (Hematoxylin & Eosin)

For the Hematoxylin & Eosin histological analysis, the samples were immersed in increasing concentrations of ethanol (70%-80%-90%) for 10 minutes each, then twice in isoparaffin for 60 minutes each, and twice in paraffin for 60 minutes, in order to embed them in a paraffin block and prepare them for microtome sectioning into 4-5µm slices and further standard H&E staining protocols.

#### 2.3.5 Immunofluorescence

For the immunohistochemistry-immunofluorescence (IHC-IF) assays, thin-slice samples were incubated in 1 µg/mL mouse anti-Sox2 primary antibody (sc-365823, Santa Cruz Biotechnology, TX, USA) in a shaker at 4°C overnight. Samples were washed 3 times with PBT 0.1% to remove excess unbound antibodies and then incubated in a mixture of 3.3 μg/mL of AlexaFluor 488 anti-mouse secondary antibody (ThermoFisher Scientific, Waltham, USA) and DAPI in a shaker at RT for 2 hours. Samples were then washed again, mounted on glass microscope slides and cover-slipped with Dako fluorescence mounting medium (S302380-2, Agilent, CA, USA).

These IHC-IF steps were carried out after incubation in R1 and H2O2 bleaching but before incubation in TDE, due to the restrictions of the clearing process.

For the lectin vasculature labeling, animals were injected intravenously through the tail with 0.3 ml of 0.52 μmol/L 649-lectin stock (DL-1178, Vector Laboratories, USA) and left in circulation for 5 minutes. They were then transcardially perfused with 5 ml of 10 U/ml heparin in PBS 1×, followed by 10 ml of 1% PFA. A further 10mL of lectin were perfused for 3 minutes, followed by 30 mL of 4% PFA. Organs were then dissected, post-fixed and cut in half before being subjected to optical tissue clearing protocols.

### 2.4 Sample property measurements

#### 2.4.1 Transparency

The effect of the optical tissue clearing process on the samples was studied by calculating their attenuation coefficient (µ) throughout the different stages of the clearing protocol. A higher attenuation coefficient corresponds to higher sample opacity and to lower light penetration.

The attenuation coefficient was calculated from the Beer-Lambert law equation:

Which results in:

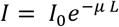

Where:

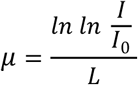

● ***I*** is the intensity of the light being transmitted through the sample.
● ***I0*** is the intensity of the incident light.
● ***L*** is the thickness of the sample being traversed by the light.

Bright-field microscopy images of the samples were acquired to obtain the light intensity value ***I***, as well as images of empty glass slides with no samples of them (keeping the same device parameters as the sample images) to obtain the incident light intensity ***I0***. The value of ***L*** was measured using precision calipers at the imaging point on each sample.

The calculations were carried out through a MATLAB® script. 3 different images were acquired for each sample, and 5 different measurements were carried out on each image; the resulting measurements were all averaged to obtain a single average µ value for each sample.

#### 2.4.2 Fluorescence intensity

The intensity of the specific fluorescence of both endogenous proteins (GFP, tdTomato) and exogenous fluorescent molecules (DAPI, AlexaFluor antibody-bound probes) was measured through the signal-to-background (SBR) ratio of fluorescence microscopy images of the samples.

The calculations were implemented as part of a MATLAB® script. This script applies a median filter to despeckle the image and applies a top hat filter to remove the background. It then segments the fluorescence signal by means of Otsu or Triangle methods and applies a combination of closing and opening morphological operations along with an area filter to remove small, noisy components, returning the average of the fluorescence signal. Autofluorescence is quantified as the mean intensity of the regions surrounding each detected fluorophore. The size of the neighborhood is given by the standard deviation of a gaussian filter, which is applied to the fluorescence image to diffuse each fluorophore and obtain its surrounding neighborhood. The resulting output is a single value for the ratio between the fluorophore’s signal and the background noise/autofluorescence. Higher SBR values correlate with higher specific signal, whereas values approaching 1 mean little or no specific signal (e.g., due to fluorescence quenching or increase in background noise).

#### 2.4.3 Volume

To measure sample volume before and after the clearing process had taken place, 3D CT scans of the different organ samples were carried out using a Molecubes X-Cube CT (Molecubes NV, Gent, Belgium) and volume was measured from the resulting image stack using the Fiji-ImageJ image processing software.

### 2.5 Imaging techniques

#### 2.5.1 Bright-field microscopy images

Both transparency measurement images and H&E images were acquired with a Nikon Eclipse E800 (Nikon, Minato, Tokyo, Japan) bright-field microscope, with light coming from below, with a Nikon Plan UW 2× objective (NA: 0.06, WD: 7.5 mm) (Nikon, Minato, Tokyo, Japan) connected to a Nikon DXM1200F digital camera (Nikon, Minato, Tokyo, Japan) mounted above the sample.

Additional images were acquired using an Olympus BX51 reflected light microscope (Olympus Scientific Solutions, Tokyo, Japan) with an Olympus MPlan 50X objective.

#### 2.5.2 Confocal microscopy images

High-resolution fluorescence microscopy images of the samples were acquired using a Leica TCS SPE inverted Confocal Microscope (Leica Microsystems, Germany) from the Confocal Microscopy Research Support Service (SAI) of the Hospital’s Experimental Medicine and Surgery Unit (UMCE). Both ASC APO 10×/0.30 DRY and ACS APO 20×/0.60/IMM objectives were used for this purpose; the 20× objective required glycerol immersion of the glass slides carrying the samples.

#### 2.5.3 SPIM images

3D Selective-Plane Illumination Microscopy (SPIM) images of the cleared samples were acquired using a custom-built SPIM device. This system uses mirrors and lenses to convert a laser beam of specific wavelength to a light sheet and guide it towards the sample (suspended in immersion media inside a glass cuvette), illuminating a single plane at a time for acquisition with an objective placed perpendicular to the plane.

The laser light sheet is focused onto the focal plane by an infinity-corrected 5× long working distance objective (NA: 0.14, WD: 34 mm, depth of focus (DF): 14 μm) (Mitutoyo Corporation, Japan). Image acquisition was carried out with a 2× objective (NA: 0.055, WD: 34 mm and DF: 91 μm) (Mitutoyo Corporation, Japan) and a 5× objective (NA: 0.055, WD: 34 mm and DF: 91 μm) (Mitutoyo Corporation, Japan) connected to a Neo 5.5 sCMOS camera (Andor, UK) with 2560 × 2160 active pixels and a physical pixel detector size of 6.5μm × 6.5μm.

A more detailed overview of the SPIM system can be found in (Nehrhoff et al. 2016).

### 2.6 Image processing

The images obtained during this project were examined and processed using the Fiji package based on the ImageJ Open-Source image processing software (Schindelin et al. 2012). 3D volumetric rendering of SPIM-acquired image stacks was carried out with the 3DSlicer software (Fedorov et al. 2012).

## 3. Results and Discussion

### 3.1 TDE can be used as a RI-matching reagent as part of an effective optical tissue clearing method (WaC)

Discerning the type of pigments or otherwise light-blocking substances to be removed is critical for the development of a new tissue clearing protocol, as universal applicability is a highly regarded feature. For this preliminary purpose, major histological structures of interest were roughly ordered from most to least pigmented according to both the amount and abundance of the biochromes and phenomena conferring them color, and the observable coloration per se (**Table 2**).

**Table 2:** Various organs ordered from most to least pigmented, along with their respective biological pigments and the color they most commonly yield.

| ORGAN/TISSUE | MOST ABUNDANT PIGMENTS | RESULTING COLOR |
| --- | --- | --- |
| LIVER | Cytochromes, hemoglobin, bilirubin, carotenes | Dark red, dark brown |
| SPLEEN | Hemoglobin, cytochromes, hemosiderin | Dark red, yellow |
| KIDNEY | Cytochromes, hemoglobin | Pink, brown |
| STRIATED MUSCLE | Myoglobin, hemoglobin, cytochromes, structural color* | Red, pink |
| BONE MARROW | Hemoglobin | Red, brown |
| HEART | Myoglobin, cytochromes, hemoglobin, carotenes | Red, brown |
| EYE | Eumelanin, pheomelanin, carotenes | Brown, black, various iris colors |
| SKIN | Eumelanin, pheomelanin, carotenes | Black, brown, yellow, white |
| BONE | Collagen | White |
| BRAIN | Myelin | Beige, pink |
| PANCREAS | Carotenes | Pink, yellow |
| BLADDER | Intrinsic color | Pink, white |
| STOMACH, INTESTINES | Intrinsic color | Pink, white |

We first wanted to evaluate whether TDE could be successfully applied as a refractive index homogenizing reagent for akin to Reagent 2 in the standard CUBIC protocol, to take advantage of the benefits the former presents over other reagents with a similar function. For this purpose, organs were subjected to a R1-incubation step and then incubated with TDE 97%. Control samples were subjected to the CUBIC protocol to compare the results.

Both methods proved to be effective at rendering the samples reasonably transparent, with TDE showcasing similar-or-better results than CUBIC (**Figure 1**). No side effects or significant downsides or failures were observed, therefore confirming our hypothesis, and verifying that TDE can indeed act as a main RI-matching agent on several organs in combination with a lipid extraction process.

**Figure 1:**
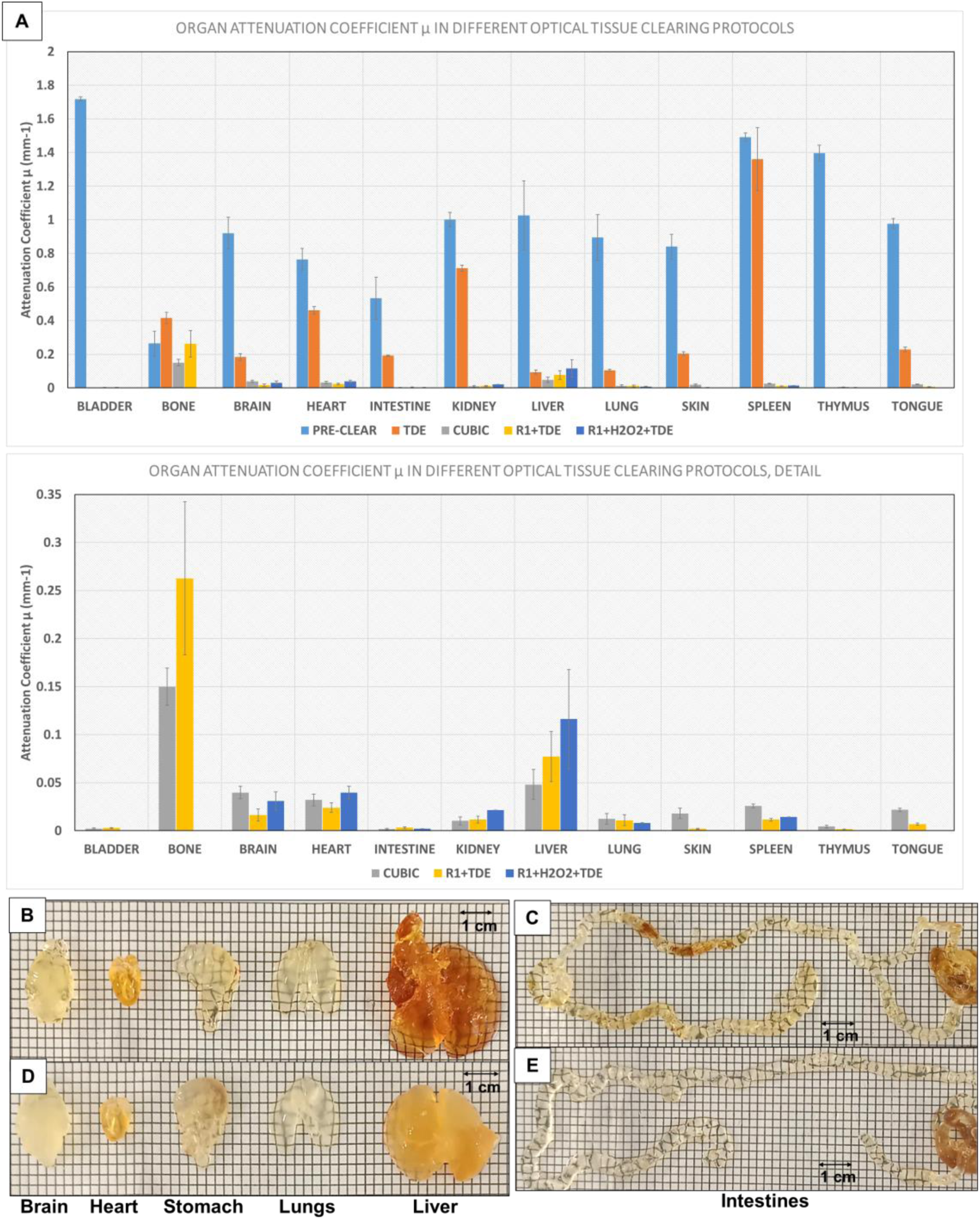
Effect of TDE, R1+TDE and R1+H2O2+TDE clearing protocol on whole mouse organs compared to CUBIC. **A)** Clearing efficiency of TDE, R1+TDE and R1+H2O2+TDE in whole mouse organs compared to CUBIC. Value for µ (Attenuation coefficient, mm^-1^) for each sample after clearing. Lower values mean higher transparency and more effective clearing action. N=5 for CUBIC and R1+TDE, N=3 for TDE and R1+H2O2+TDE. Error bars given as standard error. **B,C,D,E)** Results of clearing with **B,C)** CUBIC and **D,E)** R1+H2O2+TDE CUBIC: 10 days R1 + 1 day R2, 37°C, shaker. Bleaching: 45min 10% H2O2 in PBS, 65°C, glass container. R1+ TDE: 10 days R1 + 1 day TDE 97%, 37°C, shaker.

TDE was also tested on its own to check whether it could act as a single-step clearing protocol for certain organs; while opacity was reduced in all studied samples, this reduction proved insufficient for full-scale 3D imaging and could not compete with the results of other established methods for whole organs, despite the reduction in protocol length and increase in ease of application (**Figure 1**). Refraction Index matching alone does not seem to be enough to provide full transparency on most large-scale samples, therefore requiring extra steps to remove lipids and pigments.

Further assays to check light penetration into samples cleared with only TDE as well as R1+TDE (White and Clear, WaC) showed the same results, with TDE alone increasing the maximum light penetration depth but not achieving enough transparency to noticeably enhance clearness in most organs compared to the full protocol with the lipid removal step (**Figure 2**).

**Figure 2:**
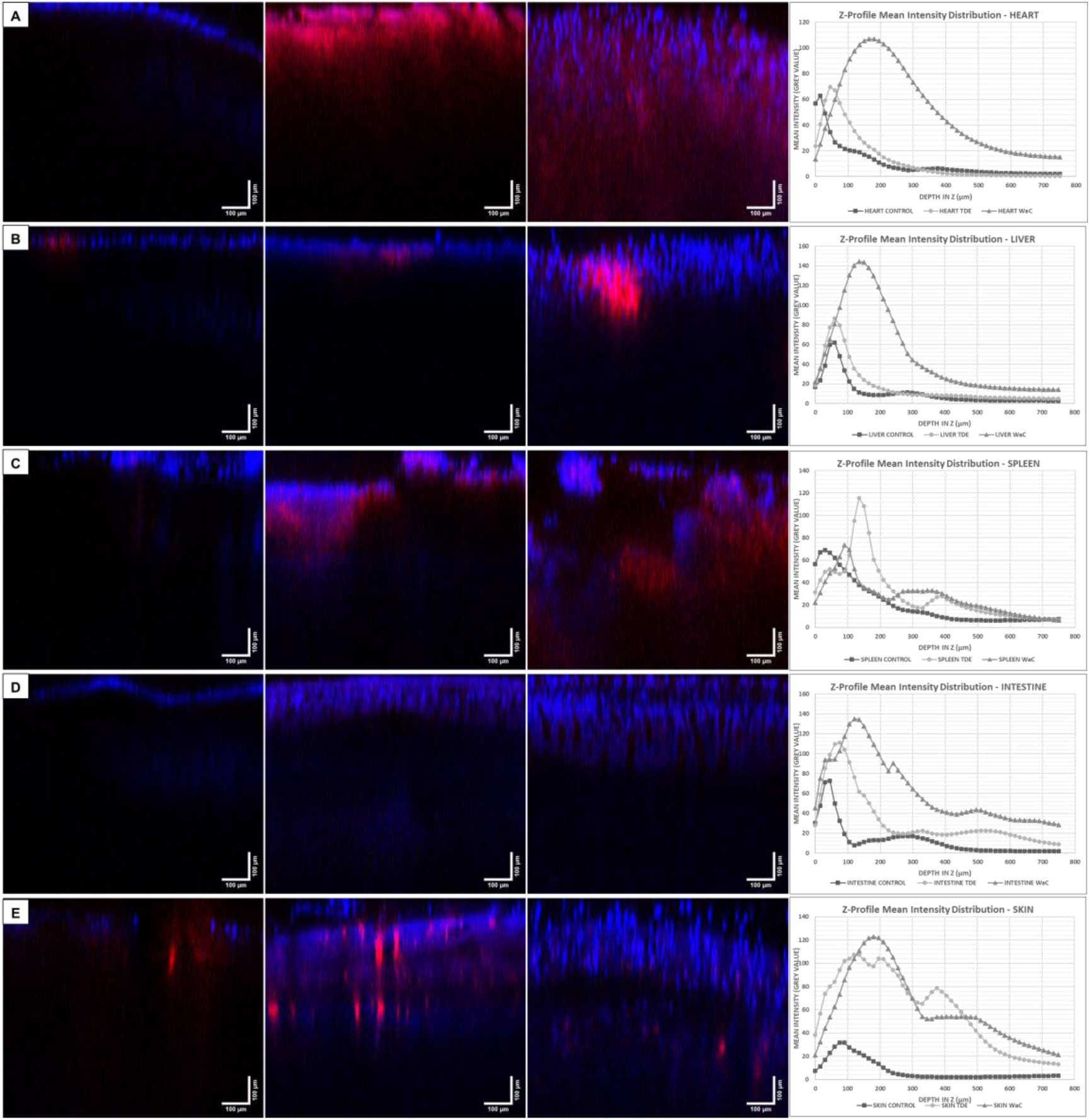
Laser light penetration in thick (∼750µm-1mm) mouse organ samples. Images obtained from uncleared samples (left), samples cleared with TDE (center), samples cleared with White and Clear (right), and Z-profile mean intensity distribution in DAPI. 10x objective, Stack depth: 750µm. Blue: DAPI (cell nuclei), Red: Lectin (vasculature). **A)** heart, **B)** liver, **C)** spleen, **D)** intestine, **E)** skin.

As part of the improved optical tissue clearing method, a new transcardial perfusion protocol was devised, where both Reagent 1 and hydrogen peroxide were perfused after blood clearance and fixation. The use of perfusion as the delivery mechanism alongside conventional immersion of samples or organs allows optical tissue clearing reagents to better penetrate tissue and bleach pigments such as the heme chromophore from hemoglobin, speeding up the clearing process and increasing its efficiency. Another variant of this protocol only included the R1 step, to rule out any possible detrimental effects of H2O2 application while verifying those of R1 perfusion itself and the ensuing optical clearing carried out by TDE.

As a result, optical clearing and pigment removal was significantly sped up, with organs presenting a noticeable increase in both when compared after only 36h incubation in R1and 12h incubation in TDE; this change was most pronounced in certain organs like the heart and spleen (**Figure 3**).

**Figure 3:**
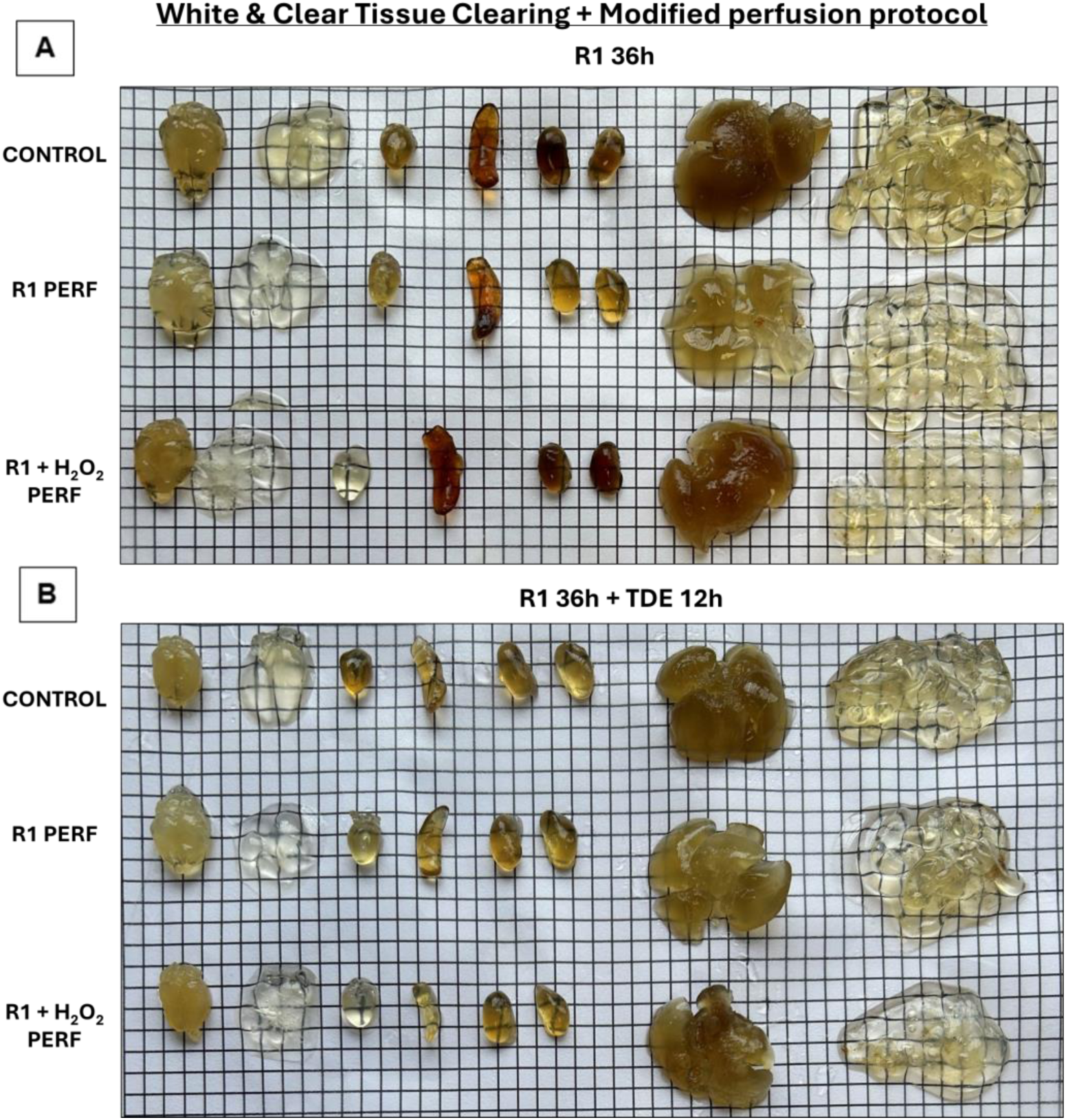
Results of optimized perfusion protocol for White & Clear tissue clearing. **A)** Whole mouse organs after 36h incubation in R1 post-perfusion using the modified perfusion protocol. 1] Control (no perfusion), 2] R1 perfusion, 3] R1 + H2O2 perfusion. **B)** Whole mouse organs after 36h incubation in R1 and 12h incubation in TDE 80% post-perfusion. 1] Control (no perfusion), 2] R1 perfusion, 3] R1 + H2O2 perfusion.

### 3.2 TDE can temporarily reduce fluorescence intensity for endogenous fluorescent proteins in a concentration- and sample-dependent manner

TDE has been reported to have a negative effect on organic fluorescent proteins such as GFP; thus, we wanted to check whether that was the case for our TDE-enhanced protocol as well. We incubated heart and liver tissue samples from transgenic mice with endogenous fluorescent proteins (TdTomato and EGFP) in different concentrations of TDE, to verify this quenching effect and test whether it was concentration dependent.

As previously described, we could observe a noticeable decrease in signal intensity relative to background (Signal-to-Background Ratio, SBR) with time for samples incubated in 100% TDE. After 24h, it became difficult to identify fluorescence with some samples seemingly losing all signal altogether (**Figure 4 A-E**).

**Figure 4:**
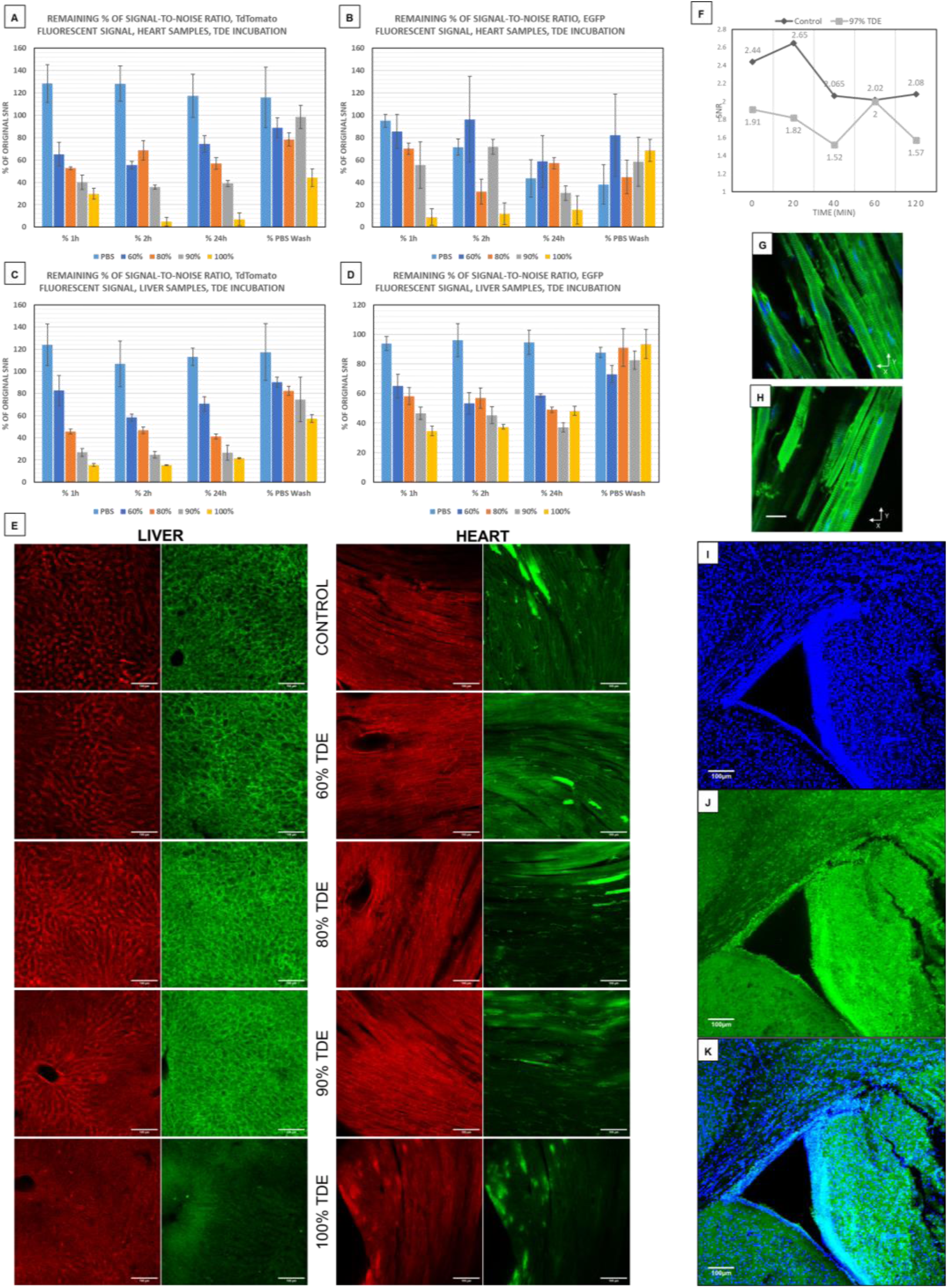
Effect of TDE incubation in endogenous TdTomato and EGFP fluorescence in LysMCre^+/−^, mT/mG mouse heart samples and AlbCre^+/−^, mT/mG mouse liver samples. **A, B, C, D)** % of the original Signal-to-noise/background ratio values over 24h for TdTomato in samples incubated in PBS (control), TDE 60%, 80%, 90%, and 100% plus additional post- incubation PBS wash for **A)** TdTomato and **B)** EGFP in heart samples, **C)** TdTomato and **D)** EGFP in liver samples. N=3. Error bars shown as standard error. **E)** Representative images of TdTomato (left, red) and EGFP (right, green) visualization in liver samples after 24h incubation in PBS, 60% TDE 80% TDE, 90% TDE and TDE 100%. Confocal microscopy images, 10x magnification. F,G,H) Effect of TDE incubation in AlexaFluor 488 fluorescence. F) Signal-to- noise ratio values over time for AlexaFluor 488 in samples incubated in PBS (control) and TDE 97%. G,H) examples of DAPI (Blue) and AlexaFluor 488 (Green) visualization in samples incubated for 1h in B) PBS (control) and C) TDE 97%. 20x Confocal Microscopy images. Scale bar = 37.2µm. I,J,K) Results of DAPI (Blue) and Sox2 (Green) IHC labelling in R1+H2O2+TDE- cleared mouse brain samples. 10x magnification confocal microscopy. I) DAPI, B) Sox2, K) channel merge. Bleaching: 45min 10% H2O2 in PBS, 65°C, glass container. R1+ TDE: 10 days R1 + 1-day TDE 97%, 37°C, shaker.

Decreasing concentrations of TDE showed a noticeable reduction in the amount of quenching experienced by the endogenous fluorescence, however, with the effect being minor in concentrations at or below 80% and enough SBR being preserved to allow visualization of the labeled tissues and structures (**Figure 4 A-E**).

Interestingly enough, the effect of TDE on endogenous fluorescence seemed to vary between different types of tissues as well as between fluorophores, as the same experiment carried out on two organs yielded different results. In the case of liver samples, while higher concentrations of TDE continued to result in the most noticeable loss of fluorescence, these changes were less pronounced than in heart tissue, with SBR remaining at more manageable levels throughout, including the maintenance of some amount of signal at 100% concentration in the case of EGFP.

This loss of fluorescence seemed to be reversible as well, as washing the samples in PBS for 1h after TDE incubation resulted in fluorescence being visible again even for samples incubated in 100% TDE which had seemingly lost all signal; this recovery was consistent across both fluorophores and organs, although the amount of recovered signal varied across the different conditions.

We then moved on to testing whether it also applied to fluorescent dyes and probes used in IHC/IF assays, as any negative impact on those would hinder the use of this modified clearing protocol for research and sample analysis.

Mouse heart samples were cut into 200µm-thick slices and a IHC protocol was applied to label cell nuclei with the DAPI fluorescent dye and α-actin present in the Z-disc of sarcomeres in cardiomyocytes with the AlexaFluor 488 IHC/IF probe; samples were then incubated in PBS and 97% TDE for 2 hours. No significant reductions in fluorescent signal were observed, and SBR values remained consistent throughout; DAPI visualization was unaltered as well (**Figure 4 F-H**).

Thus, we concluded that TDE does not seem to have any negative effects on signal from fluorescent dyes or commercial AlexaFluor IHC/IF probes.

Next, we wanted to check whether the combination of clearing and bleaching in the same protocol could influence the immunoreactivity of target proteins. Mouse brain samples were cleared with the full WaC protocol alongside an intermediate IHC step to label cell nuclei with the DAPI fluorescent dye and Sox2 (a neural stem cell and neural progenitor cell marker) with the AlexaFluor 488 IHC/IF probe, and subsequently sliced into 1mm-thick pieces to acquire confocal microscopy images.

The IHC assay was successful, with Sox2+ cells being correctly labeled, and their presence found in relatively high numbers in one of the main NSC niches, the subventricular zone (SVZ). DAPI nuclear labeling was successful as well, with no major alterations or disturbances found throughout the whole sample (**Figure 4 I-K**).

We thus believe immunoreactivity and structure are indeed preserved by the modified clearing and bleaching protocol, and thus IHC/IF assays performed on samples cleared using this protocol would be valid for the purposes of any study or structural or functional analysis.

### 3.3 WaC has no significant effects on tissue and organ properties compared to other clearing methods

CUBIC can have an engorging effect on certain samples, causing them to swell and increase in size; these changes are not considered to have an impact on results obtained from analysis of those samples, but are nevertheless a deviation from the original tissue’s morphology. We wanted to check whether our modified clearing protocol would have any other impact on sample size, structure and properties that could negatively impact any possible results.

All the samples displayed the expected swelling during the R1 incubation, with a resulting increase in volume compared to the original. We verified that TDE does cause certain tissues to shrink as previously reported (Aoyagi et al. 2015), although this effect is not nearly as strong as the swelling impact of R1 on those same samples (**Figure 5 A,B**). In our opinion, the reason for this reduction is the following: TDÉs clearing dynamics are governed by Fick’s first law of diffusion, with larger concentration gradients between cells and the surrounding solution speeding up TDÉs diffusion into the cell. However, they also speed up the outflow of water from the cells; as water is less dense and can diffuse more easily, these larger concentration gradients extract water faster than TDE itself diffuse into the cells. Thus, by the time an equilibrium is reached, a larger volume of solvent has left the cell than has entered it, causing a decrease in volume.

**Figure 5:**
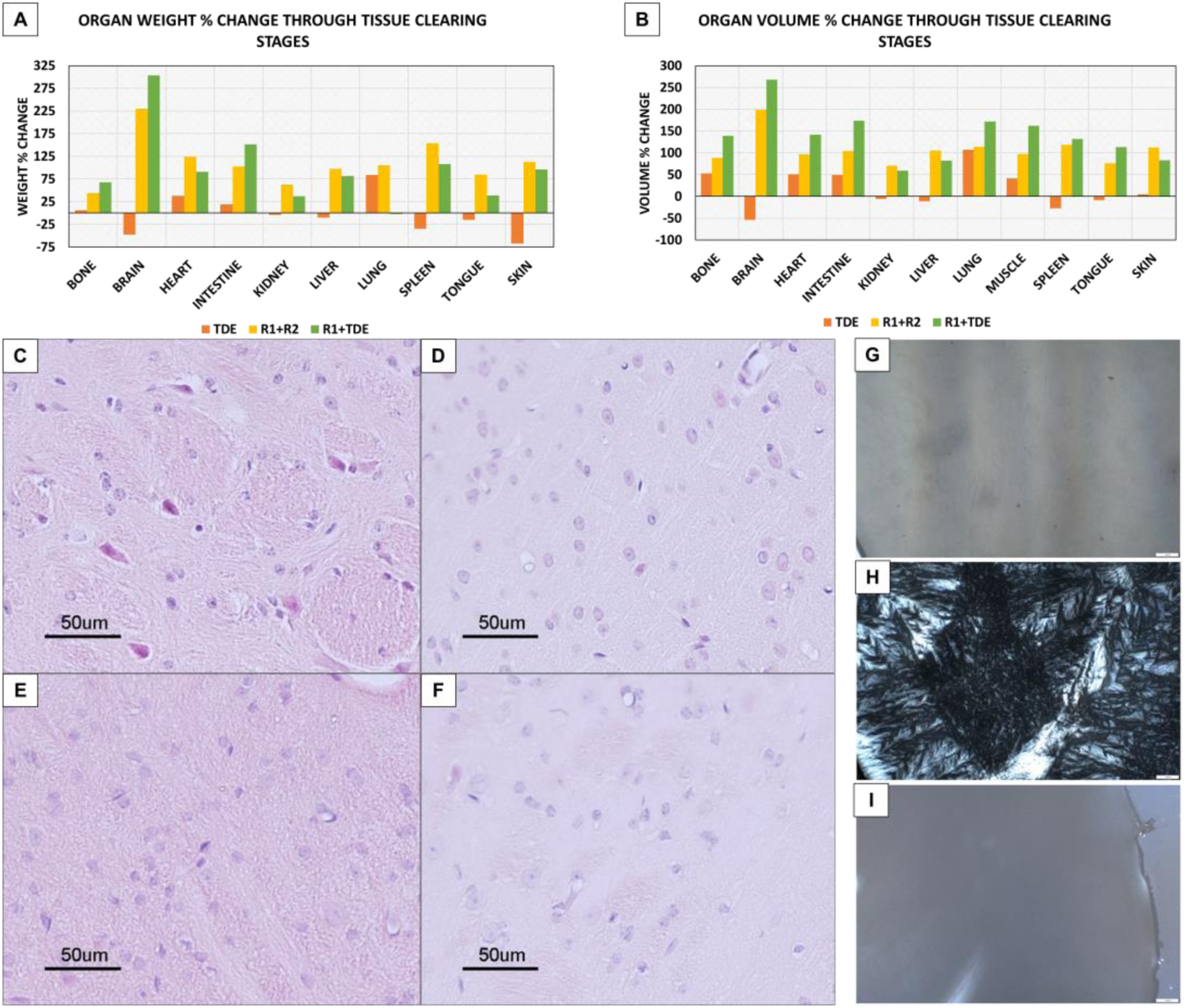
Effect of tissue clearing protocols on physical properties of the organs. **A)** Representative values for volume % change through the different stages of the tissue clearing protocols. Higher values mean a more drastic change in volume. **B)** Representative values for weight % change through the different stages of the tissue clearing protocols. Higher values mean a more drastic change in weight. **C,D,E,F)** Hematoxylin & Eosin studies of cleared brain samples. **C)** Control, **D)** R1+TDE, **E)** R1+H2O2+TDE, **F)** H2O2+R1+TDE. 20x Bright-field microscopy. Bleaching: 45min 10% H2O2 in PBS, 65°C, glass container. R1+ TDE: 10 days R1 + 1-day TDE 85%, 37°C, shaker. **G,H,I)** Bright-field microscopy images of **G)** control, **H)** CUBIC-cleared and **I)** WaC-cleared thin mouse brain slices showcasing leftover crystal precipitations in samples in R2 and no change in samples in TDE. 20x magnification.

It should be noted that organs in which a further increase in volume or weight was observed after R1+TDE incubation did not correspond to those were TDE alone caused an increase in the first place; while the trend of R1-induced swelling seems consistent across all organs, the specific magnitude of this effect as well as whether TDE itself causes swelling or shrinking seems to show great variability across the same organs from different animals.

We also carried out conventional Haematoxylin & Eosin histological studies on the cleared brain samples, to verify the lack of any significant alterations to tissue structure or morphology that could negatively affect IHC assays and analysis. Upon examination, no significant alterations were observed (**Figure 5** C-F). The tissue seemed to remain unaltered between the samples and when compared to the control images, with the only difference in visualization being the intensity of the dyes; all samples incubated in R1 showcased a loss of intensity, while those treated with H2O2 after R1 seemed to recover part of the expected tint.

These results confirmed previous claims of preservation of tissue morphology throughout the H2O2 bleaching protocol, as well as the lack of negative effects of TDE on the same properties.

Close examination of cleared samples also showed the accumulation of crystals in samples incubated in R2, presumably created by precipitation of the sucrose that makes up Reagent 2; samples incubated in TDE showed no traces or alterations compared to the control, further reinforcing the preservation of sample composition and structure showcased by TDE (**Figure 5 G-I**).

The use of TDE as an RI-matching alternative is desirable due to its properties and the ensuing advantages it offers over other existing clearing reagents, such as CUBIC’s own Reagent 2; the only aspect in which TDE shows a slight disadvantage by comparison is its effect on endogenous fluorescence, but even then this can be mitigated through chosen concentration values (**Table 3**).

**Table 3:**
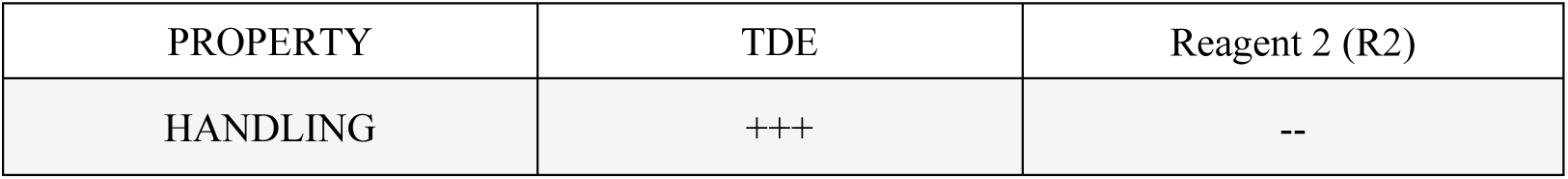
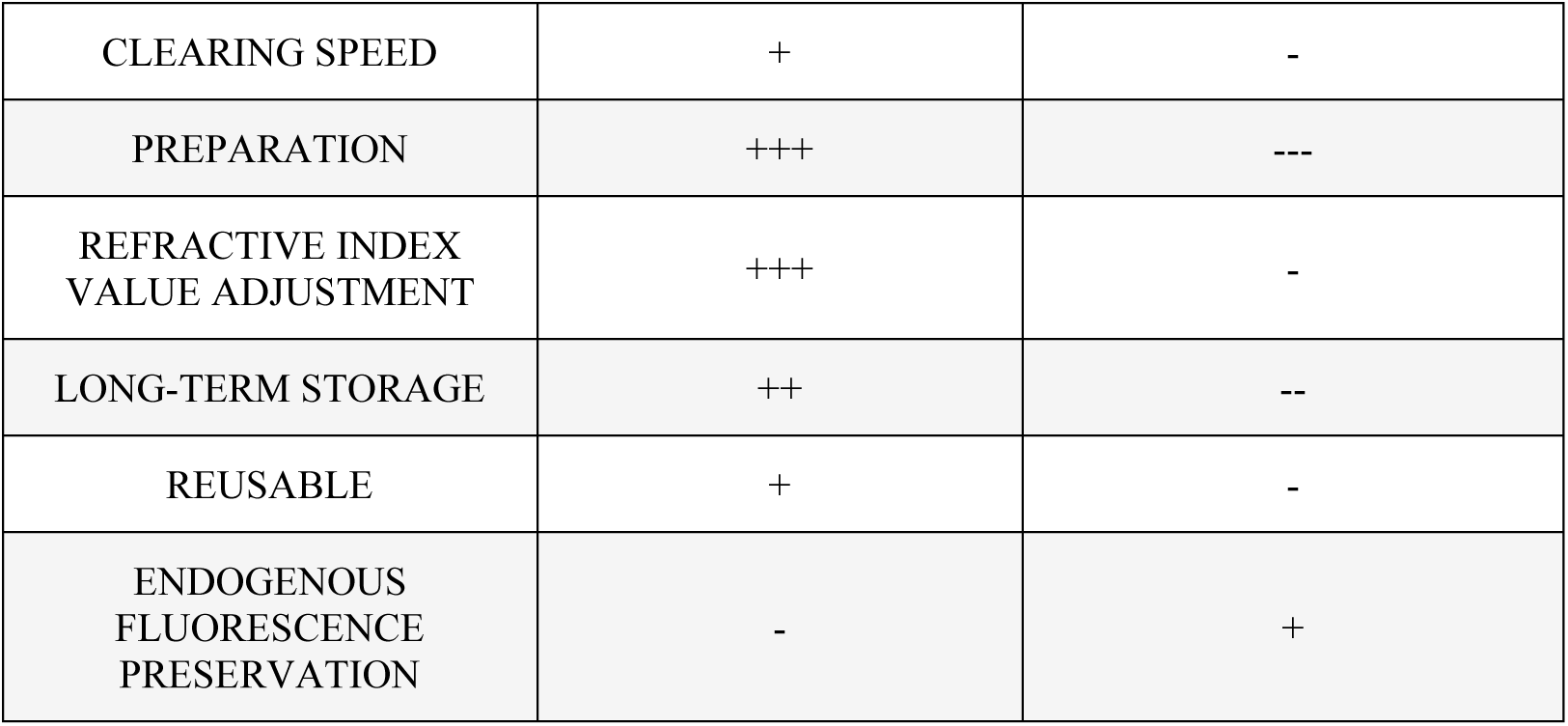
Comparison between select properties of TDE and CUBIC’s Refractive Index matching Reagent 2.

| PROPERTY | TDE | Reagent 2 (R2) |
| --- | --- | --- |
| HANDLING | +++ | -- |
| CLEARING SPEED | + | - |
| PREPARATION | +++ | --- |
| REFRACTIVE INDEX<br>VALUE ADJUSTMENT | +++ | - |
| LONG-TERM STORAGE | ++ | -- |
| REUSABLE | + | - |
| ENDOGENOUS<br>FLUORESCENCE<br>PRESERVATION | - | + |

The main advantage showcased by TDE in this case is its reduced viscosity compared to R2, which both increases the speed of reagent penetration into the samples (and thus the clearing speed) and makes it possible to implement injection and removal of TDE through automated or mechanical systems, which would otherwise be prone to clogging or low flow speeds.

Another advantage of TDE over similar reagents is the possibility to adjust its RI value by changing the concentration, allowing to tailor the specific RI values to different tissues and samples while maximizing the resulting transparency and/or fine-tuning the value for each imaging system’s own optimal RI requirements. In order to verify whether specific organs favored certain concentrations, several organs were cleared using W&C and then incubated in different TDE concentrations, with the attenuation coefficient being measured and compared between them (**Figure 6**). Due to it seemingly providing the best results across most organs, as well as reducing the impact on endogenous fluorescent proteins (if any), 85% was chosen as the default for W&C, with other concentrations being adjustable for certain organs depending on whether endogenous fluorescence is present and on which concentration is best suited for maximum transparency in each organ, e.g. brains favoring higher concentrations (97%-100%) instead for absolute maximum efficiency.

**Figure 6:**
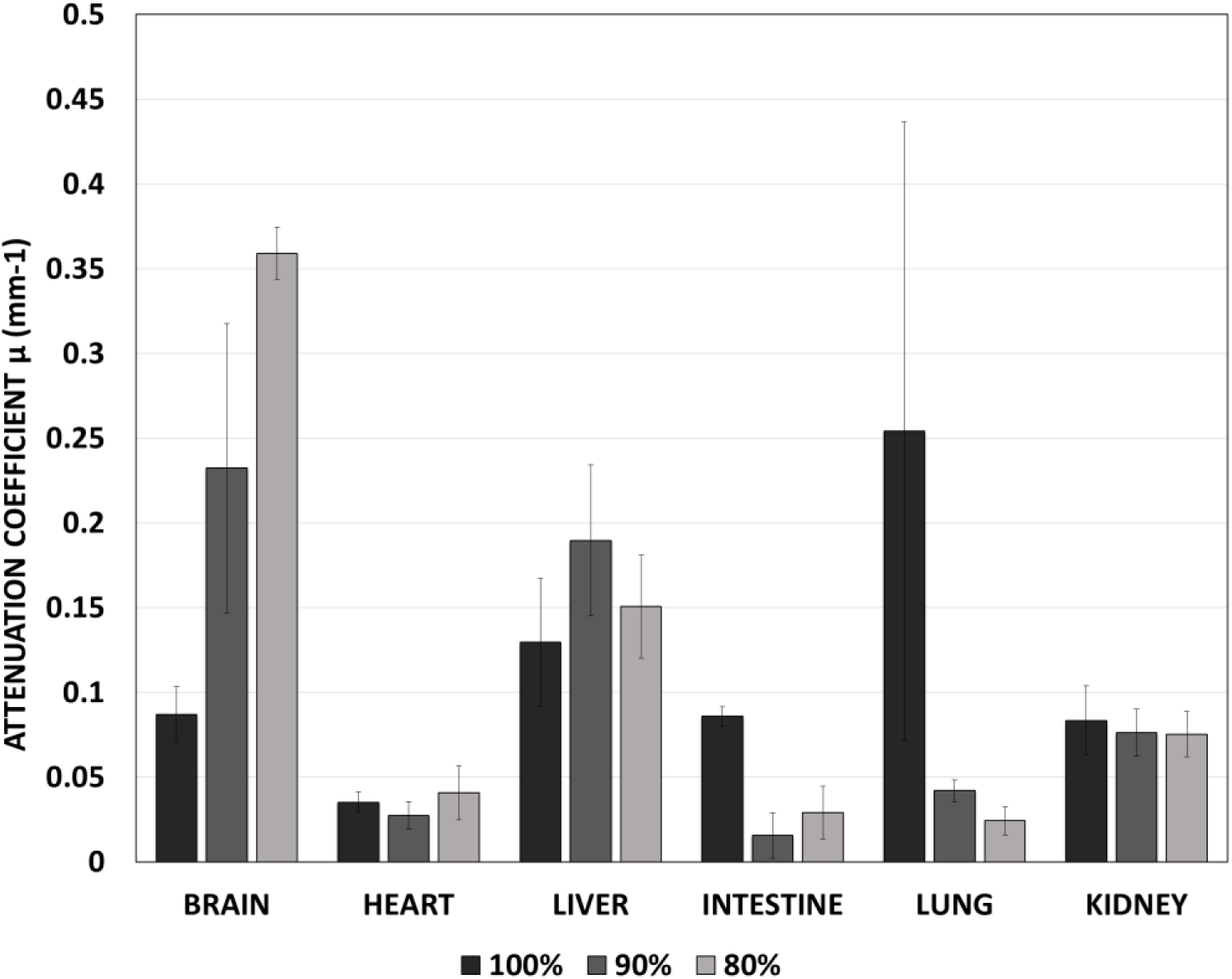
Attenuation coefficient values for W&C-cleared mouse organs incubated in different TDE concentrations. N=3 for each concentration and organ. Error bars given as standard error.

Last, but not least, preparation and storage of TDE is a very easy process, as it is a single commercially available reagent with a long shelf life which only needs to be diluted to the target concentration and does not precipitate when kept at low temperatures and does not require any complex preparation steps nor any special conservation procedures.

### 3.4 WaC can be used in combination with IHC/IF methods to label and clear thick slices and whole-organ samples

We moved on to attempting an optical tissue clearing and IHC/IF labeling procedure on larger-scale samples, to check whether there would be any effects on antibody penetration or performance and verify this tissue clearing protocol was compatible with these applications. This assay was successful as well, with brain samples being properly labelled to detect the mature neuron marker NeuN and the vascular endothelial marker CD31 (**Figure 7 A,B**).

**Figure 7:**
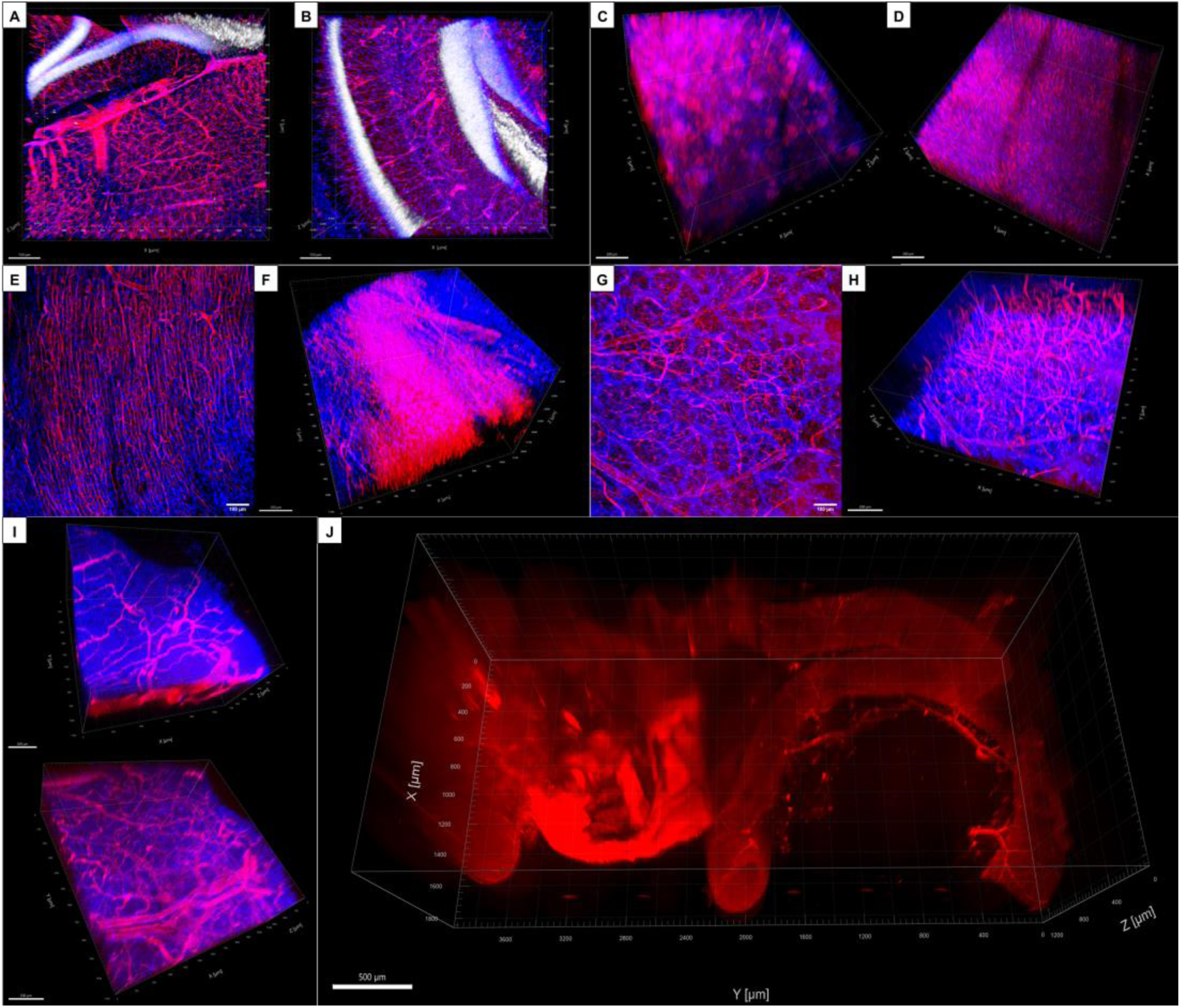
Results of WaC clearing and labeling on different organs and samples. **A, B)** IHC/IF labelling on WaC+H2O2-cleared thick mouse brain samples. Blue: DAPI (cell nuclei), Red: CD31 (vasculature), White: NeuN (neurons). **A)** 10x confocal microscopy, Slice depth: 3,5mm, Stack depth: 230µm. **B)** 10x confocal microscopy, Slice depth: 4mm, Stack depth: 255µm. **C,D)** Lectin labeling on WaC-cleared whole mice organs. Blue: DAPI (cell nuclei), Red: Lectin-labeled vasculature. 10x magnification confocal microscopy. **C)** 3D reconstruction of vascular network on the kidney (stack depth: 500µm). **D)** 3D reconstruction of vascular network on the liver (stack depth: 400µm). **E,F)** Lectin labeling on WaC-cleared whole mice hearts. Blue: DAPI (cell nuclei), Red: Lectin-labeled vasculature. 10x magnification confocal microscopy. **E)** Representative 2D image of vascular network on the left ventricle wall (Z-projection, max intensity), **F)** 3D reconstruction from complete image stack (stack depth: 720µm). **G,H)** Lectin labeling on WaC-cleared whole mice brains. Blue: DAPI (cell nuclei), Red: Lectin-labeled vasculature. 10x magnification confocal microscopy. **G)** Representative 2D image of vascular network on the left olfactory bulb (Z-projection, max intensity), **H)** 3D reconstructions from complete image stack (stack depth: 500µm). **I)** Lectin labeling on WaC-cleared whole mice stomach. Blue: DAPI (cell nuclei), Red: Lectin-labeled vasculature. 10x magnification confocal microscopy, (stack depths: 700µm/400µm). **J)** Lectin labeling on WaC-cleared whole mice intestine. Red: Lectin-labeled vasculature. 2x SPIM microscopy.

As a next step, we applied a lectin-based vasculature labelling method in order to image the vascular network and check the potential interactions with the tissue clearing protocol. For this purpose, we applied a reduced R1+TDE clearing protocol to acquire large-size image stacks corresponding to large sample volumes. Hearts from animals subjected to the labelling protocol were immersed in R1 for 3 days and subsequently washed and immersed in 85% TDE o.n. This assay was successful again, with the vasculature being clearly labeled and visible up to remarkable depths through the tissue, and with no evidence of ill-effects on the labelling (**Figure 7 E-J**). In the case of certain organs such as the intestine, it was possible to acquire whole-organ 3D images with the use of a SPIM microscope due to the level of transparency achieved by the clearing protocol.

We thus have further reasons to state that immunoreactivity and structure are indeed preserved by the modified clearing and bleaching protocol, and thus IHC/IF assays performed on samples cleared using this protocol would be valid for the purposes of any study or structural or functional analysis.

## 4. Conclusions

By taking biological pigments and time optimization into special consideration, a new tissue clearing protocol that bypasses rarely addressed complications was successfully brought forth, namely W&C. Firstly, the drawbacks caused by CUBIC’s standard RI-matching reagent, Reagent 2, such as a considerably time-consuming preparation due to the highly viscous nature of its components in solution (which also results in a poorer, slower penetration into the samples to be assessed), are effectively corrected by employing TDE instead; ready-to-use and free-flowing, not only are its handling easier and performance enhanced when compared to Reagent 2, but the issue of crystalline byproducts precipitating onto samples upon incubation due to the high concentrations of sucrose in the makeup of Reagent 2 is completely avoided. Its considerably lower viscosity also brings the possibility of automatization closer, as the risk of clogging pumps or nozzles is dramatically decreased. Furthermore, by diluting TDE beyond its unadultered presentation, its RI-matching properties can be readily customized depending on the organ to be cleared and, ultimately, the protocol’s users’ needs.

Although Reagent 2 does retain the advantageous feature of organic protein-borne fluorescent retention over TDE, the signal loss in the latter is easily reversible upon an overnight PBS wash, even in cases where any traces thereof appear to be completely missing. Signal recovery is also consistent across all tested types of tissue samples and sources of endogenous fluorescence.

Other aqueous-based clearing methods’ delipidation step is fundamental in the removal of pigments and intrinsic coloration that renders organs opaque (in other words, solely relying on RI-matching does not provide satisfactory results); however, non-lipophilic pigments, particularly hemoproteins, normally remain intact. W&C rids organs of the deep red that may stubbornly hang on to highly vascularized and mitochondria-rich organs due to, respectively, leftover hemoglobin and cytochromes through a so-called bleaching agent: diluted hydrogen peroxide introduced through both perfusion and incubation. The use of hydrogen peroxide may elicit concerns over the physical integrity of the treated sample; through a simple H&E analysis it was ascertained that, save for some tinction loss, all histological structures remain otherwise unperturbed.

As W&C displays an exceptionally versatile performance, it is evident that the development of an improved clearing protocol that is also universally applicable to all organs was adequately achieved.

